# Dissecting the heat stress altering immune responses and skin microbiota in fish in a recirculating aquaculture system in Singapore

**DOI:** 10.1101/2024.01.02.573918

**Authors:** Tze Hann Ng, M Sobana, Xian Zhe Chew, Thiviya Nair D/O Madhaven, Jing Wen Chow, Adrian Low, Henning Seedorf, Giana Bastos Gomes

**Author notes:** Address correspondence to Giana Bastos Gomes. Thiviya Nair D/O Madhaven, Danish Shellfish Centre, National Institute of Aquatic Resources, Technical University of Denmark, Denmark. Adrian Low, Department of Medicine, Yong Loo Lin School of Medicine, National University of Singapore, Singapore.

## Abstract

Environmental factors, probiotics and feed additives affect microbiota diversity in fish. Water temperature disrupts the composition and diversity of microbiota, with temperature changes triggering stress and immune responses in fish. In Singapore, water heat treatment is used to control and prevent disease outbreaks in farmed fish. Although gut microbiota responses to heat stress have been reported, little is known about the effects of heat treatment on fish immune responses and fish skin microbiota dynamics. Over a 3-mo interval, we investigated microbiota dynamics and fish immune responses associated with a heat treatment practice in a commercial fish farm with a recirculating aquaculture system (RAS). Tank water temperature was raised to 37-39 °C for 1 hour, every 2 weeks. Tank water and fish microbial communities were analysed by 16S amplicon sequencing, and host molecular expressions were determined by qPCR. We inferred that heat treatment increased stress and immune responses that protected fish from opportunistic infections. However, overreaction to temperature stress can cause dysbiosis of the skin microbiota and death. We also suggested the value of the skin microbiota Pseudomonadota: Bacteroidota (P:B) ratio as a biomarker for aquaculture fish health.

**IMPORTANCE:** Aquaculture is an emerging economic activity to supply high-quality animal protein and reduce reliance on wild-caught fish products. Recently, there has been emphasis on intensive aquaculture, using a Recirculating Aquaculture System (RAS). In RAS, management of pathogens/parasites prevalence is a major challenge. Developing practical solutions for producing healthy juveniles in nursery systems will make profound contributions to sustainable aquaculture. In this study, we used an unconventional strategy, exposing juveniles to the pathobiome in the environment, followed by non-lethal heat shock treatments to enhance immunity. Short-term stress induced heat shock proteins that protected fish from opportunistic infections. We concluded that manipulating environmental-microbial-host interactions, together with enhanced functional capacity of fish immune response, has potential for disease control in aquaculture.

## INTRODUCTION

Commensal and opportunistic bacteria in fish skin mucus are part of fish skin microbiota (1). The fish primary immune response is extremely active in mucus (2, 3), providing protection against environmental pathogens. However, many pathogenic and opportunistic bacteria can adhere to skin mucus, evade the fish defence barrier, and promote infection; overgrowth of these bacteria can disrupt the relative abundance of commensal bacteria (dysbiosis). However, not all skin microbial dysbiosis causes disease (4–8).

Gut microbiota have been evaluated in >145 species of teleosts (9), whereas there has been limited assessment of skin microbiota of farmed fish species (10–12). Skin microbiota can be influenced by many factors including fish species and geographic origin (13, 14). Early research (15) demonstrated seasonal changes in skin bacterial abundances in North Sea cod. Fish in warmer waters have more mesophiles, whereas fish near coasts have more halotolerant bacteria (15, 16). The microbiota of unstressed fish is dominated by taxa with probiotic and antibacterial properties, whereas the microbiota of fish under stress seems to be dominated by potential pathogens (17). Crowded conditions and low oxygen stress fish and cause dysbiosis in the skin microbiota, promoting growth of opportunistic pathogens (17–19). Furthermore, diet, acidification and salinity variation can also contribute to microbial dysbiosis in fish (20).

Current climate changes, including warming water, alters the equilibrium of the fish microbiota (20, 21). Water temperature disrupts the composition and diversity of the gut and skin microbiota of healthy fish. The equilibrium of skin and gut microbiota were disturbed by temperature fluctuations in chum salmon, especially *Vibrio* in fecal samples (20). During warm seasons, growth of potentially pathogenic *Vibrio* species may be promoted by water temperature. However, many *Vibrio* species are hardly pathogenic, but mainly opportunistic when changes in water quality parameters occur, particularly in controlled aquaculture systems (22, 23). Temperature changes can increase bacterial metabolism, allowing production of virulent genes that can promote infection and disease outbreaks (24). However, increased temperature may also have the opposite impact (25). Rhabdoviruses in salmonid and Japanese flounder, betanodaviruses in European sea bass (26), and *Yersinia*, *Flavobacterium*, *Lactococcus*, and *Vibrio* genera (24), have all been linked to temperature restriction that affects virulence in fish where reduced mortality has been observed.

Temperature fluctuations can have combinatorial effects on fish; drastic changes in temperature impacts fish, by changing the immune system and metabolic rates (27, 28). In salmon infected with parasites, water was heated to 34 °C for 30 seconds (29); sea lice separated from the host and parasite’s adult stage was reduced. In a study in Artemia larvae, a non-lethal heat treatment of 37 °C for 30 min increased resistance against *Vibrio* species known to infect brine shrimp. Induction of a stress response following heat treatment was linked to the protective effect (30). The main responsive factors during stress events are stress proteins, namely heat shock protein (Hsp) genes including Hsp70, and Hsp90 that had either downregulated or upregulated expression profiles when exposed to temperature changes (31, 32). Moreover, Hsps have a crucial role in mounting protective immune responses against infections (32, 33).

Limited research has been done on heat treatment for disease management in aquatic animals. Understanding and manipulating environmental-microbial-host interactions together with enhanced functional capacity of fish immune response could promote development of a more sustainable aquaculture. In this study, samples were collected from a commercial farm in Singapore that raises *Lates calcarifer* (barramundi) with heat treatment practices for disease management, allowing us to examine how fish immune response and skin microbiota vary with temperature. To analyze the microbiota, fish fin and kidney, biofilm, and water samples were subjected to 16S rRNA-based metagenomic analyses. Protective impacts were assessed by the state of health and immune responses in the liver. Insights from the present study could unleash key information related to stress biomarkers to be used in disease surveillance for fish health in aquaculture.

## Materials and Methods

### Fish rearing and heat treatment in a commercial fish farm

#### Overview

To study the suitability of heat treatment practices for disease management, we collaborated with a commercial fish farm in Singapore that uses a recirculating aquaculture system (RAS). This farm raises barramundi in tanks [tank size; diameter: 8.45 meter, height: 2.55 meter, volume: 143 m^3^] with a capacity of up to 12 tons of fish per tank. Fish transfers in and out of tanks are regular operations that may introduce infections. The farm uses heat treatment with water temperatures up to 37–39 °C for 1 hour, aiming to eliminate pathogens (especially viruses) and prevent disease outbreaks.

#### Sampling

Sampling was done for 13 weeks (20 Sep 2021 to 13 Dec 2021) with weekly water and biofilm collected from six tanks. Two liters of water were sampled from each tank and transported on ice to the Temasek Life Sciences Laboratory in Singapore for laboratory processing. Each water sample was filtered via a 3.0-µm, 47 mm-diameter (Merck, US) membrane, followed by another 0.22-µm, 47 mm-diameter (Whatman, Germany) membrane to recover microbial cells. Membrane filter papers swabs of biofilm from the tank wall were put into 2.0-ml Eppendorf tubes and stored at -20 °C until further processing. However, some water and biofilm samples could not be collected due to practical and safety issues at the commercial farm.

Both Healthy (n=6) and Moribund (n=5) fish (based on their appearance and clinical signs) were removed from one of the heat-treated tanks on two heat treatment days, whereas Control (n=15) fish were sampled on three non-heat treatment days. Fish were immediately euthanized with Aqui-S® (40 mg/L; AQUI-S, New Zealand), length and weight recorded, and then placed on ice. The sampled fish ranged in size from 12.5-20.7 cm. Samples of brain, liver, kidney, spleen, and fin were collected from each fish and preserved in 95% ethanol at - 80 °C pending DNA/RNA extraction.

### Microbiological compositions by 16S-based sequencing

#### DNA extraction and 16S rRNA gene amplicon sequencing

A total of 73 water and 20 biofilm samples were used for DNA extraction based on a CTAB/chloroform-isoamyl alcohol protocol (modified from 34). Whereas 26 fin and 16 kidney samples were extracted using the DNeasy Blood & Tissue Kits (Qiagen) according to the manufacturer’s instructions, including pretreatment for Gram-positive bacteria with Enzymatic lysis buffer (20 mM Tris·Cl, pH 8.0; 2 mM sodium EDTA; 1.2% Triton® X-100; 20 mg/ml lysozyme) at 37 °C one hour and proteinase K digestion for three hours. The DNA samples were sent to NovogeneAIT Genomics Singapore Pte. Ltd. for quality control, library construction and sequencing. PCR amplification of V4-V5 regions of the 16S rRNA gene was done using specific primers linked to barcodes. PCR products were confirmed using 2% agarose gel electrophoresis. The library was examined using a Qubit and bioanalyzer for size distribution identification and quantification. Equal amounts of PCR products from each sample were pooled and further ligated with Illumina adapters. To produce 250 bp paired-end raw reads, libraries were sequenced on a paired-end Illumina NovaSeq platform (Illumina, USA).

#### Data processing

Libraries (n=135) were constructed and sequenced for the V4–V5 regions. Based on the unique barcode, paired-end reads were assigned to their respective samples. Quality filtering on the raw reads was done to obtain high-quality clean reads. Barcode and primer sequence were trimmed and reads < 60 bp removed. Based on the read overlap, paired-end clean reads were combined using FLASH (V1.2.7). Chimera was detected using UCHIME and removed to obtain nochimera reads.

Data processing of the sequencing reads was performed using the pipelines in QIIME2 Version 2021.4 (35). Briefly, reads were quality-filtered and denoised using DADA2 (36). Taxonomy was assigned using the SILVA pre-trained classifier (silva-138-99-nb-classifier); each unique taxonomic features represents an amplicon sequence variant (ASV). All chloroplast, mitochondria or unidentified sequences, and ASVs that were less frequent than 15 or observed in less than three samples were removed. Alpha diversity (Shannon diversity) and beta diversity (Bray–Curtis dissimilarity) were calculated in Phyloseq (37) using an even depth across samples (rarefaction depth of 90% of the minimum sample depth, in this study 34, 826 was chosen). Taxonomic summary at phylum and genus levels were used for further analysis on taxa with a relative abundance >1%. The top ten taxa were sorted by abundance based on total abundance across all samples using top_taxa function (R package “microbiome”; 38). Statistically significant variation in microbiota composition and diversity was detected by analysis of similarity (ANOSIM) and pairwise Wilcoxon tests.

### Host genes quantification

#### RNA extraction and Quantitative Polymerase Chain Reaction (qPCR)

Total RNA was extracted from liver samples using the RNeasy Plus Micro Kit (Qiagen) and Invitrogen SuperScript IV - First Strand Synthesis Kit (Invitrogen) was used to synthesize complementary DNA (cDNA) according to the manufacturer’s instructions. Based on published sequences, primer sequences for *Hsp70*, *IL1β*, *TNFα* and *EF1α* genes were synthesized (Table 1). Internal references were provided by the housekeeping gene *EF1α*. Quantitative polymerase chain reaction (qPCR) was performed using a CFX96 (Bio-Rad). Reactions were carried out in a 20 µl mixture of 1 µl diluted cDNA, 10 µl SYBR green master, 0.4 µl forward primer, 0.4 µl reverse primer, and 8.2 µl water. The qPCR program was: initiation at 95°C for 3 minutes, denaturation at 95°C for 3 seconds, followed by 40 cycles of annealing and amplification at appropriate melting temperature for 20 seconds. The 2^-ΔCt^ method was used to calculate relative mRNA expression (39). R version 4.2.2 and R studio (40) was used for statistical analyses (R package “rstatix”; 41) and graph generation (R package “ggplot2”; 42). All data were shown as the IQR and pairwise Wilcoxon tests were used to compare means. The Benjamini-Hochberg (BH; 43) method was used to adjust the p-value for multiple hypothesis tests.

**Table 1:**
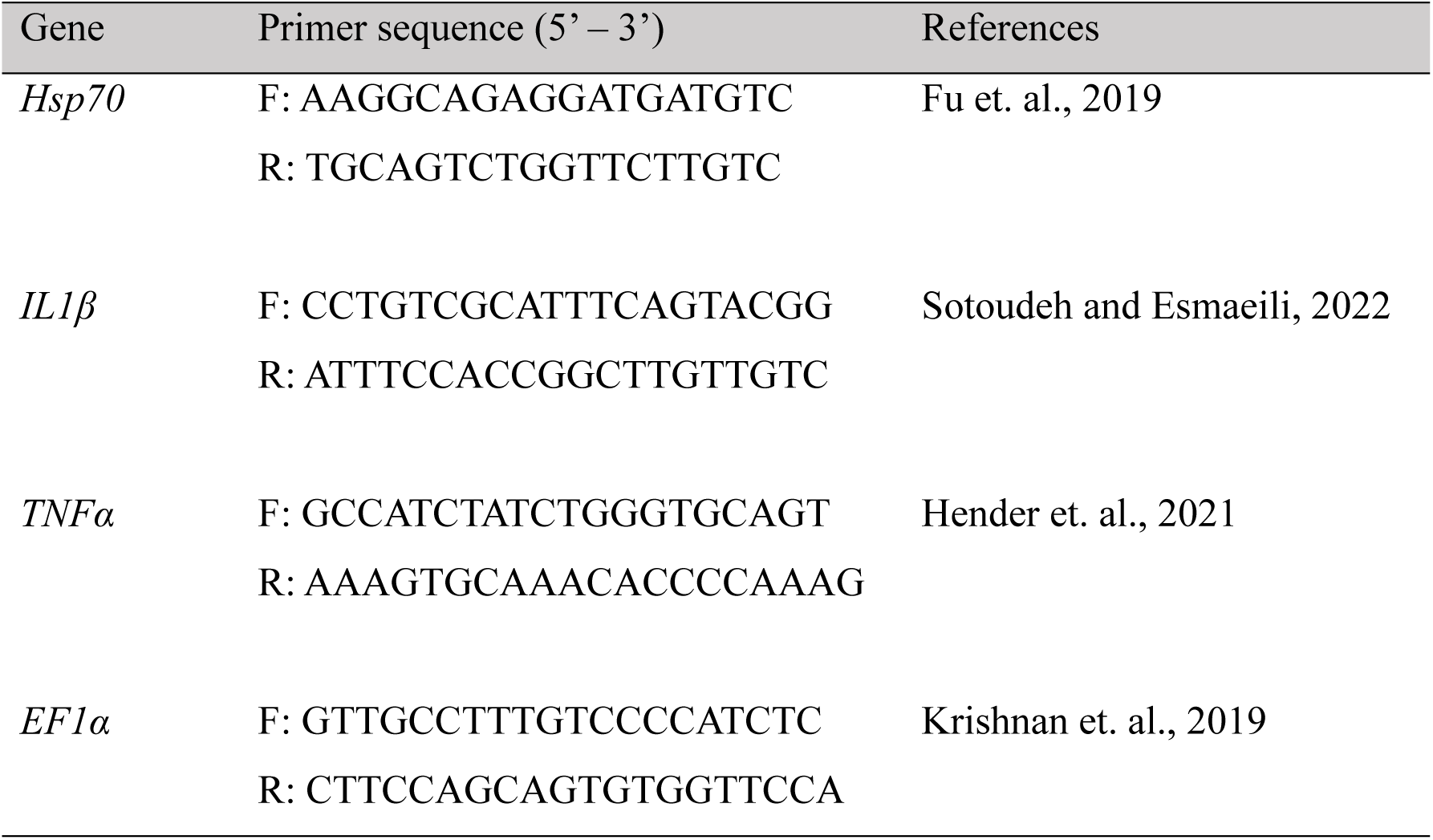
Primers used in this study.

## Results

### Sampling and sequence data

In total, 73 water bottles and 20 biofilm samples and 26 fish fin samples and 16 fish kidney samples were collected and sequenced. Approximately 26 million raw reads were retrieved; the number of reads per sample ranged from 99,917 to 289,162. These sequences corresponded to 5,355 unique ASVs, and after filtering and removing ASVs belonging to organelles, resulted in 4,802 unique ASVs for downstream analysis.

### Comparison of the environment microbial communities to fish tissues microbiota

The microbiota of fish tissues (fin and kidney) differed from those of the environment (tank water and biofilm), which formed well-separated clusters with no overlap, based on nonmetric multidimensional scaling (NMDS) ordination analysis with Bray-Curtis distance (Figure 1A). The significance of differences in sample types was then determined using ANOSIM tests (R= 0.7899, *p*=0.001). Environment sample communities had higher alpha diversity values compared to fish tissue microbiota (Kruskal-Wallis test, *p*=2.2 x 10^-16^; Figure 1B). When environmental and biofilm communities were compared, alpha diversity in biofilm was greater than water communities (*p*<0.005). Kidney had the lowest α-diversity among the sample types and was substantially lower than fin in terms of tissue microbiota (*p*<0.005). For microbial composition, Bacteroidota (formerly Bacteroidetes) and Pseudomonadota (formerly Proteobacteria) were prevalent in the environment, whereas Pseudomonadota dominated the microbiota of fish tissues (Figure 1C). The phyla Bdellovibrionota, Chloroflexi, and Myxococcota were detected only in biofilm communities. Furthermore, Firmicutes, which are typically prevalent in gut microbiota (44, 45), were <1% in fin microbiota.

**Figure 1:**
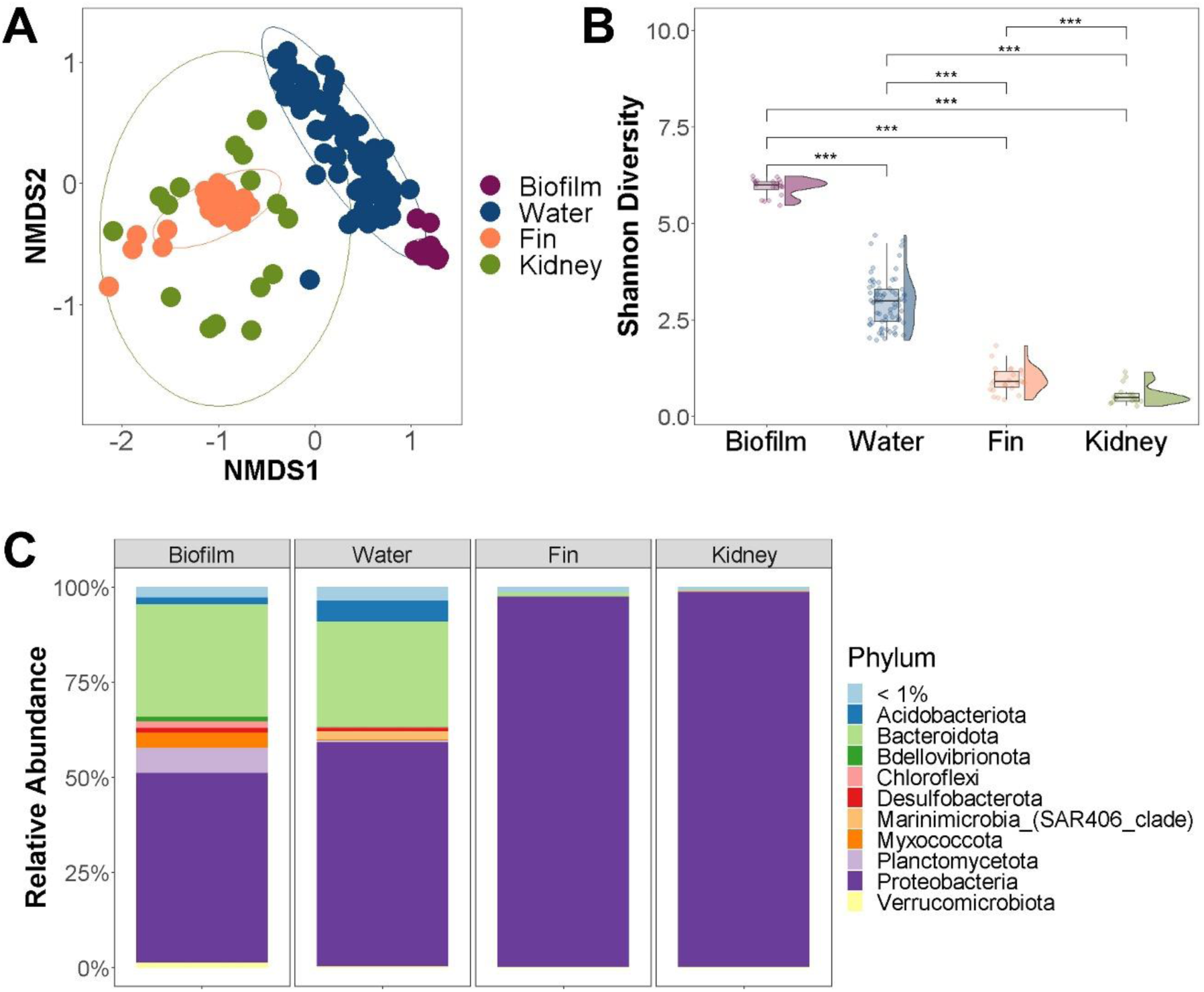
Environment bacterial communities and fish tissues microbiota. (A) Bacterial profiles were compared by non-metric multidimensional scaling (NMDS) using the Bray-Curtis distance metric displaying the community dissimilarities (based on ASVs) among sample types. ANOSIM statistic R= 0.7899, *p*=0.001. (B) Diversity plot of bacterial diversity comparing sample types based on Shannon index. Differences between sample types were tested by Wilcoxon test, pairwise comparisons *p*-value (*** *p*<0.005). (C) Relative abundance of bacterial phyla among sample types. Rare taxa in each group are indicated as < 1%.

### Effect of elevated temperature on microbiota of fish tissues

Heat treatment lowered alpha diversity values in healthy fish fin and kidney microbiota (Figure 2A). There was higher alpha diversity in the microbiota of moribund fish and were higher (*p<*0.05) in the kidney of moribund versus healthy fish. In the NMDS, fin (Figure 2B) and kidney (Figure 2C) microbiota had control tissue microbiota clustered separately from tissue samples following heat treatment (healthy and moribund fish. However, only the kidney microbiota had significant clustering (ANOSIM tests, R= 0.2165, *p*=0.037). There were no differences between healthy and moribund fish for fin microbiota (ANOSIM tests, R= 0.1195, significance=0.145).

**Figure 2:**
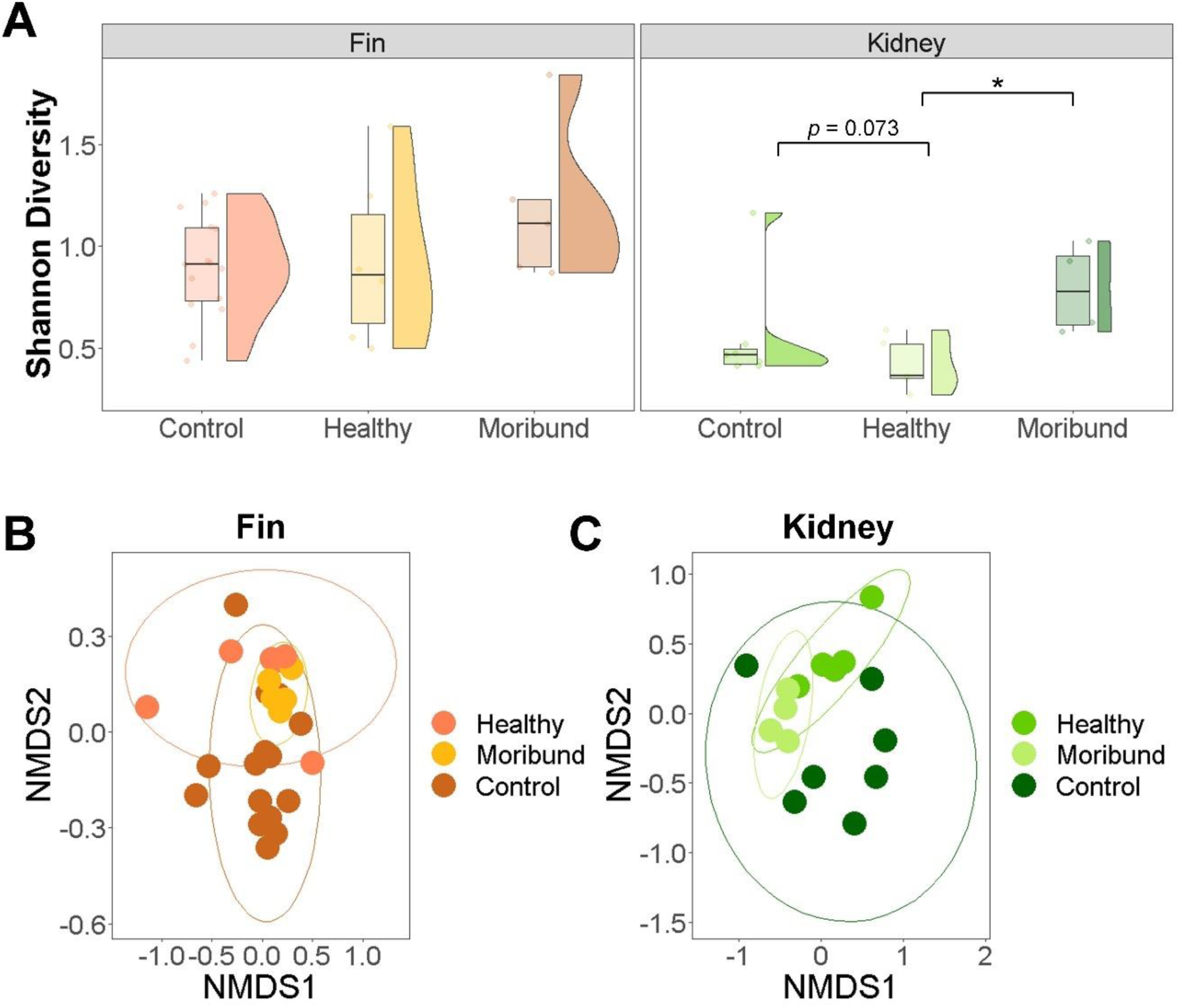
Diversity of fin and kidney microbiota in control, healthy and moribund fish in a heat-treated tank. (A) Diversity plot of bacterial diversity comparing control, healthy and moribund fish based on Shannon index. Bacterial profiles of (B) fin and (C) kidney were compared by non-metric multidimensional scaling (NMDS) using the Bray-Curtis distance metric displaying the community dissimilarities (based on ASVs) among fish tissues samples. Control: fish collected on non-heat treatment days; Healthy: healthy fish collected on heat treatment days; Moribund: moribund fish collected on heat treatment days.

### Pseudomonadota - Bacteroidota ratios reveal dysbiosis in fish tissues samples

Relative abundances of different phyla in tissue samples were analyzed, including both fin and kidney microbiota (Figure 3A). Both fin and kidney had elevated Bacteroidota relative abundances and decreased Pseudomonadota relative abundances in moribund fish (Figure 3A). As a result, the P:B ratio tended to decrease with health status; moribund fish had a lower P:B ratio (Figure 3B).

**Figure 3:**
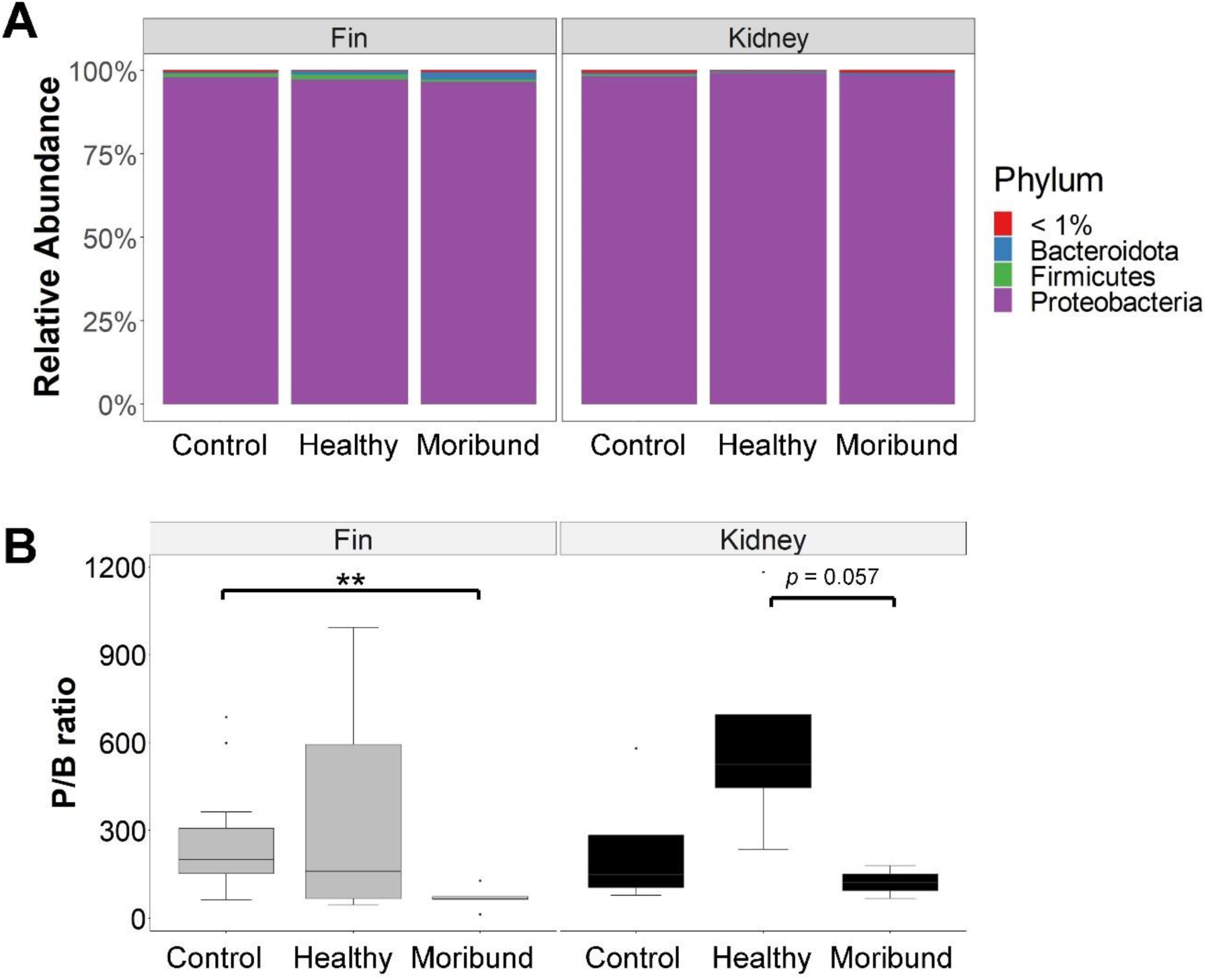
Composition of fin and kidney microbiota in control, healthy and moribund fish in a heat-treated tank. (A) Relative abundance of bacterial phyla under heat treatment. Rare taxa in each group with relative abundance < 1% indicated as < 1%. (B) P/B ratio. Differences within same sample types were tested by Wilcoxon test, pairwise comparisons significant *p*-value (* *p*<0.05, ** *p*<0.01, *** *p*<0.005) showing based on the connecting compared groups. Control: fish collected on non-heat treatment days; Healthy: healthy fish collected on heat treatment days; Moribund: moribund fish collected on heat treatment days.

### Bacteria community that contributes to dysbiosis

Relative abundances of top genera were compared among tissue samples to determine temperature effects on certain bacteria communities that contribute to dysbiosis in tissue microbiotas. The number of bacteria with significantly different relative abundance between the healthy and moribund fish microbiota at the genus level is presented in Figure 4. The genera, *Marinobacterium*, *Marivivens*, *Pseudoalteromonas*, *Pseudomonas*, *Tenacibaculum*, and *Vibrio* had significantly increased relative abundance in moribund fish. Among these genera, *Pseudomonas*, *Tenacibaculum*, and *Vibrio* are well-known fish opportunistic bacteria that can cause infection (46–48). Notably, the microbiota of both fin and kidney had a slight decrease in relative abundance of *Streptococcus*, another genus that causes illness in barramundi (49, 50).

**Figure 4:**
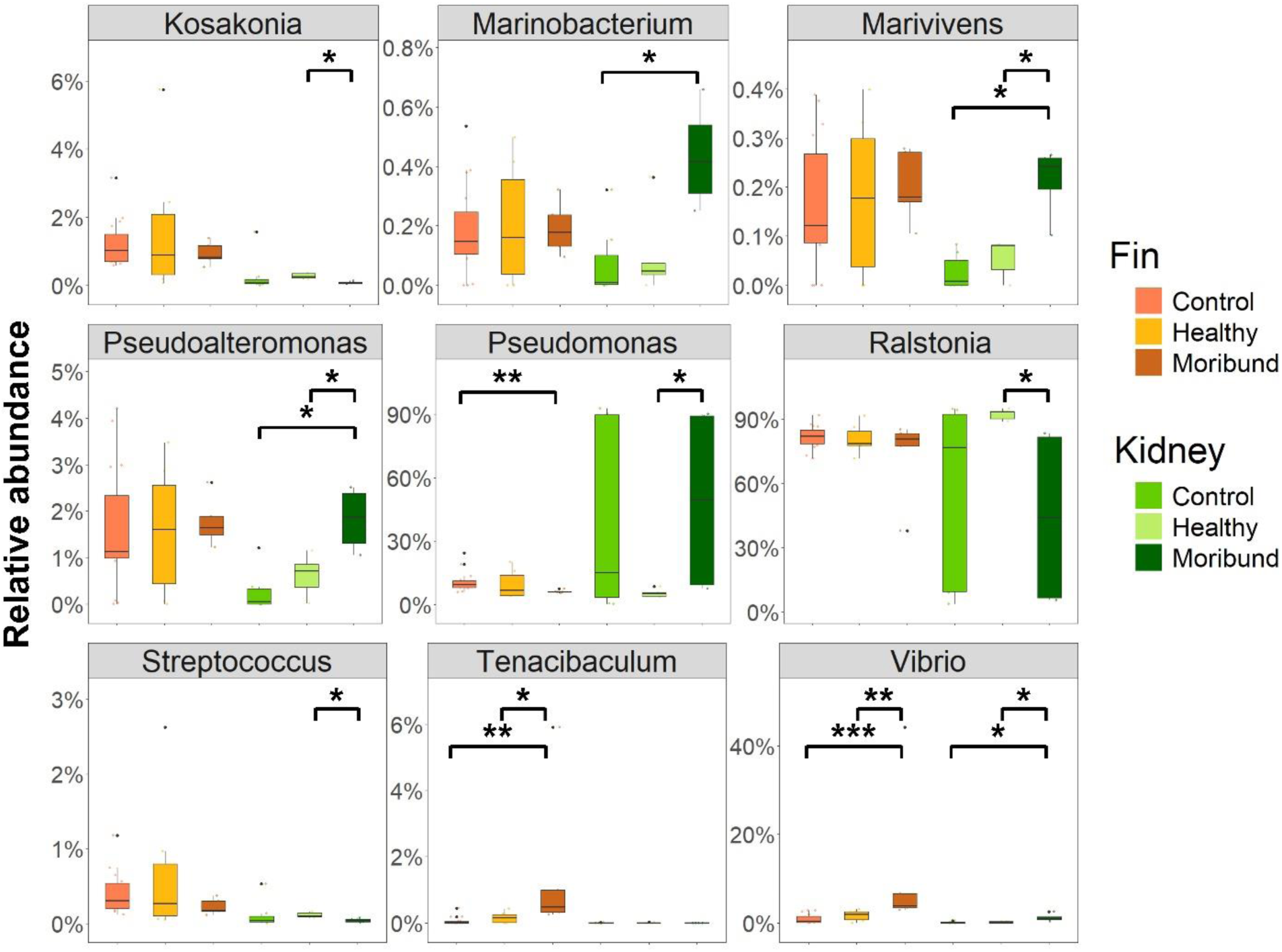
Top genera relative abundance. Differences within same sample types were tested by Wilcoxon test, pairwise comparisons significant *p*-value (* *p*<0.05, ** *p*<0.01, *** *p*<0.005) showing based on the connecting compared groups. Control: fish collected on non-heat treatment days; Healthy: healthy fish collected on heat treatment days; Moribund: moribund fish collected on heat treatment days.

### Fish immune genes response to heat treatment

The heat stress response gene, *heat shock protein 70* (*Hsp70*) mRNA level, increased dramatically as water temperature increased (Figure 5A). Moribund fish had a greater stress reaction, with a higher *Hsp70* response in moribund versus healthy fish. The proinflammatory cytokines genes, *interleukin-1-beta* (*IL1β*) and *tumor necrosis factor-alpha* (*TNFα*) had higher expression levels in healthy versus moribund fish (Figure 5B-C).

**Figure 5:**
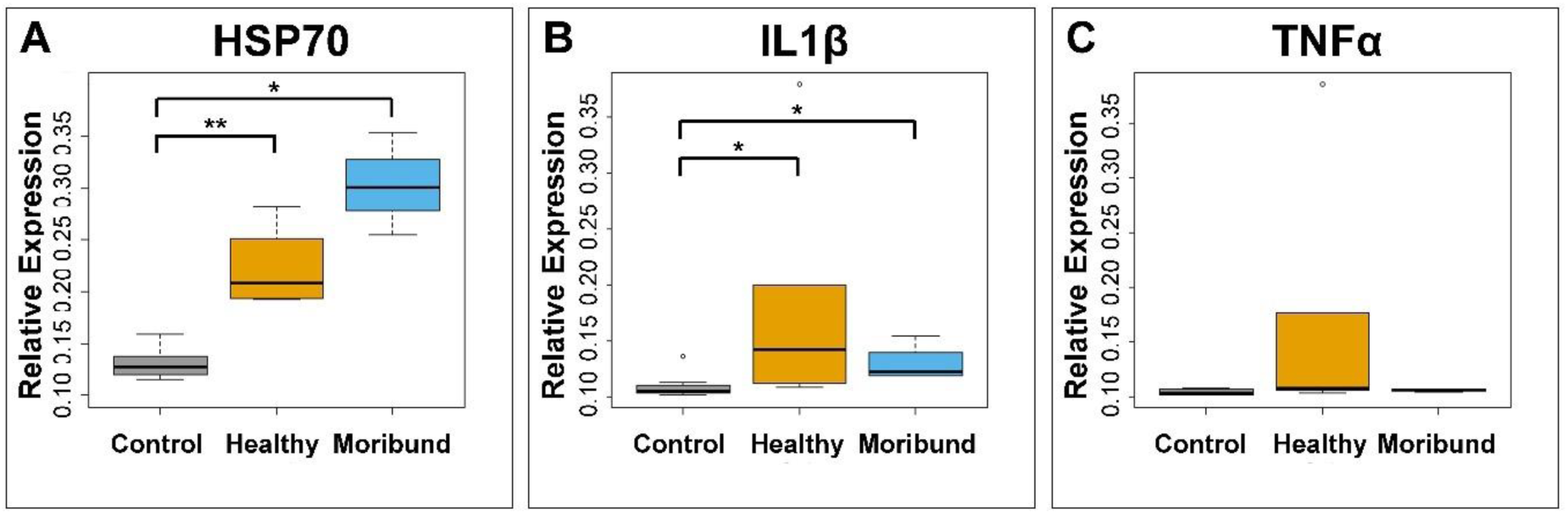
Effects of the heat treatment on host responses. Host genes (A) *Hsp70*, (B) *TNFα* and (C) *IL1β* expression was detected in the liver of barramundi using qPCR. Wilcox test was used to determine differences (* *q*<0.05, ** *q*<0.01; p.adjust.method, Benjamini-Hochberg). Control: fish collected on non-heat treatment days; Healthy: healthy fish collected on heat treatment days; Moribund: moribund fish collected on heat treatment days.

## DISCUSSION

### Temperature effect on host fitness and microbiota

The present study investigated the immune response and skin microbiota of barramundi fish subjected to water heat treatment. The host response and composition of microbiota are likely to change in response to external factors, including temperature. Water temperature changes, both increases and decreases, affect physiological performance and behaviour in fish (51), implying that cold or heat stress response is likely to affect fish health and survival. Elevated levels of heat shock proteins in fish following changes in water temperature have been reported previously (52, 53). In the farm where our study was conducted, fish were kept in tropical temperatures (range 29 to 31 °C) and heat treatment was raising the water temperature up to 39 °C for 1 hour. This increased gene expression of *Hsp70*, confirming that this heat treatment had a significant effect on the heat stress response of barramundi fish. Rainbow trout red blood cells synthesized Hsp70 in response to increased temperature and the stress protein can be induced in fish acclimated to 10 °C (54). Upregulation of Hsp70 also occurred in fish *Labeo rohita* exposed to water temperatures of 37–38 °C for 2 weeks (55).

Hsp70 is known as a “thermotolerance” protein due to its numerous and diverse contributions under heat stress conditions (56). High temperature causes protein denaturation, with Hsps conferring protection. Heat stress not only induces Hsp70, but also other heat shock protein families such as Hsp60, Hsp90, and low molecular weight Hsp30 in cell lines, primary cell culture, and also *in vivo* (31, 57). Hsps families, especially Hsp70, are not only involved in protein stabilization, but also in immune activation (58, 59). In the present study, there were increased levels of *IL1*β and *TNFα* expression in healthy barramundi following heat treatment, although the increase in *TNFα* expression was not significant. Induction of *IL1*β and *TNFα* occur concurrently in response to immune stimulation in rainbow trout (*Oncorhynchus mykiss*), Japanese flounder (*Paralichthys olivaceus*) and grass carp (*Ctenopharyngodon idella*). (60–62), but there were also observations that only *IL1*β but not *TNFα* were induced (63, 64). Similar to our findings, a possible reason is lack of an optimal sampling time point to detect significant *TNFα* upregulation, as a heat shock-induced immune response has a delayed induction of *TNFα* compared to *IL1*β (65, 66). In rainbow trout exposed to 28°C after preconditioning at 19°C for 30-60 min, there was higher immune expressions with secondary heat treatment, with upregulated *Hsp70*, *IL1β* and *TNFα* expressions (64). However, stress level of pre-conditioned fish was significantly lower than non-pre-conditioned fish. These findings implied that immunity can be modulated in fish with heat treatment, and the mechanisms in barramundi need further investigation.

Microbiota structures of the fin and kidney tissue from the present study were distinct from the structure of surrounding water microbiota, which was also the case for previous studies (67, 68). Although some bacteria derived from tissue overlapped with those of water microbiota, tissue microbiota often has specific bacterial consortia (20, 69). Bacteria composition analysis also revealed that temperature had a significant effect on diversity and taxonomic structure of the barramundi fin and kidney microbiota. In salmon gut, exposure to high water temperature favored growth of *Vibrio spp.* (69, 70). In the current study, there was a lower abundance of pathogenic bacteria *Vibrio* and *Tenacibaculum* in healthy versus moribund fish after heat treatment. In Atlantic cod, heat stress caused immune activation and limited harmful bacteria (13); bacterial degradation was detected earlier in fish kept at higher temperatures and yielded escalated expression of immune genes e.g., *IL1*β and *interferon gamma* (*IFNγ*). The authors suggested that it is unlikely that the increased temperature destroys pathogens, but rather it enables the host immune system to control the pathogens (71, 72). In our study, there was much higher *Hsp70* gene expression in moribund versus healthy fish after heat treatment. Although expression of *Hsp70* and immune genes were induced by high water temperature, it also caused fish to become weak and die. Drastic temperature shifts can cause tissue damage and mortality in aquatic animals (29, 73). Heat stress can harm fish epidermal cells and skin barrier function by reducing mucus cell numbers and disrupting mucosal microbiota (74). Therefore, in this study we inferred that heat treatment induced increases in *IL1*β and *TNFα* which controlled pathogenic bacteria and protected fish. However, in some fish, the heat treatment protocol may have induced an exaggerated response and dysbiosis that lead to death. Therefore, determining the threshold of survival temperature for heat treatment is important and may vary among fish species (51).

### Barramundi fish skin microbiota

Many current protocols to detect pathogens in farmed fish require killing animals to collect internal organ tissues (e.g., kidney) which may be undesirable for valuable or rare broodstoock (75). However, non-lethal sampling for detection of pathogens, including fin clips and mucus swabs, is growing in practice (76–78). Skin and gill microbiota can be modulated by gut health status and presumably reflect changes in fish health (11). However, limited microbiota research has been conducted on fish internal organs, and it is unclear whether they are fundamentally different from mucosal surfaces such as skin, gills, and gut lining. We propose that skin microbiota may be used to predict microbiota dysbiosis in kidney, which indicates infection status. In the present study, skin microbiota dynamics mirrored those in kidney, with Pseudomonadota dominating. There were higher abundances of fish opportunistic bacteria (e.g., *Vibrio* and *Tenacibaculum)* in skin and kidney microbiota of moribund fish, with higher significance in skin microbiota. Similar finding was also made in a previous study on farmed diseased barramundi that showed signs of tenacibaculosis (79).

To predict microbiota dysbiosis, the Pseudomonadota: Bacteroidota (P:B) ratio can be an indicator of a change in the health status of farmed fish. In the present study, there was a change in P:B ratio in healthy and moribund fish. Specific bacterial species may be effective indicators for detecting health status, but some may not correspond to the microbiota in another fish species. In addition, amplification of specific primers may have bias towards/against some taxa, making it difficult to compare studies performed with different types of primer pairs. Thus, it may be necessary to standardize the techniques (primer pairs) used when comparing taxa ratios across study. An increased Firmicutes: Bacteroidota (F:B) ratio has been linked to obesity, diabetes, ulcerative colitis and other issues in mammalians (80, 81). Firmicutes and Bacteroidota are not necessarily the major phyla of fish mucosa; in fact, Pseudomonadota tend to be much more prominent (82, 83). As a result, microbiota detection based on the ratio of P:B instead of F:B ratio could be used in fish. A previous study also proposed P:B ratio to predict fish health status (11), consistent with our observations from the present study. We observed that a change in the P:B ratio was more significant in skin samples than in kidney. We inferred that dysbiosis in fish skin communities could provide an early alert regarding opportunistic bacteria. Therefore, monitoring P:B ratio in skin microbiota by non-lethal sampling has potential to improve fish health monitoring in farmed fish.

### Limitations of the study and future research

The heat treatment under current conditions had a clearly significant impact on barramundi fish’s reaction to heat stress, as evidenced by the heat shock proteins gene expression, induced following heat treatment. After heat treatment, there were higher levels of cytokine gene expression in healthy fish, although not all genes had a statistically significant difference. Given that heat shock-induced immune responses may have a delayed induction response, and we lack sampling time points for detectable substantial effects in this study due to the operation in a commercial farm, we may have missed some important data points for a more conclusive results. Moreover, certain samples were not collected for practical and safety concerns in the farm. Therefore, it would be important to study further the impacts of water treatment on barramundi under carefully monitored settings that nonetheless closely resemble those found in a commercial production unity. Given that heat treatment may induce severe damage, more than protective effect in fish, alternative approaches should be explored. Perhaps testing different protocols of heat treatment where fish is exposed to heat for a shorter period of time, could have a more positive outcome. Further studies should also include different fish ages to understand how different fish stages of development are impacted by heat treatment. Another suggestion for further exploration, would be adding specific supplements in feed to induce heat shock response in fish (84, 85).

## ACKNOWLEDGEMENTS

We thank the collaborating farm from Singapore to allow real-time sampling.

## AUTHORS’ CONTRIBUTIONS

THN and GBG conceived the study. SM and XZC conducted the field sampling. THN performed data sorting and analyses. SM, TNM, JWC and BHL performed laboratory assessments. THN wrote the manuscript. GBG and HS reviewed and edited the manuscript.

## DATA AVAILABILITY STATEMENT

Raw sequences are available in NCBI’s Sequence Read Archive (BioProject PRJNA1055435).

## ETHICS APPROVAL

The fish collection was approved under the Institute Animal Care and Use Committee (IACUC) approved project IACUC-2019-A08 James Cook University Singapore.

